# Structure-based design of antisense oligonucleotides that inhibit SARS-CoV-2 replication

**DOI:** 10.1101/2021.08.23.457434

**Authors:** Yan Li, Gustavo Garcia, Vaithilingaraja Arumugaswami, Feng Guo

## Abstract

Antisense oligonucleotides (ASOs) are an emerging class of drugs that target RNAs. Current ASO designs strictly follow the rule of Watson-Crick base pairing along target sequences. However, RNAs often fold into structures that interfere with ASO hybridization. Here we developed a structure-based ASO design method and applied it to target severe acute respiratory syndrome coronavirus 2 (SARS-CoV-2). Our method makes sure that ASO binding is compatible with target structures in three-dimensional (3D) space by employing structural design templates. These 3D-ASOs recognize the shapes and hydrogen bonding patterns of targets via tertiary interactions, achieving enhanced affinity and specificity. We designed 3D-ASOs that bind to the frameshift stimulation element and transcription regulatory sequence of SARS-CoV-2 and identified lead ASOs that strongly inhibit viral replication in human cells. We further optimized the lead sequences and characterized structure-activity relationship. The 3D-ASO technology helps fight coronavirus disease-2019 and is broadly applicable to ASO drug development.

## INTRODUCTION

ASO drugs has the potential to revolutionize medicine ^1, 2^. ASOs hybridize to target RNAs and elicit therapeutic effects either by sterically blocking them from cellular or viral machinery or by inducing their degradation ^3^. ASO drugs are chemically modified to achieve nuclease resistance, stability in serum and cellular compartments, and favorable pharmacokinetics ^4^. The simplicity of Watson-Crick (WC) base pairing rules in principle allows any sequences to be targeted by ASOs. Hence, ASO therapeutics can regulate cellular pathways that are not readily druggable using traditional protein-binding small molecules ^5^. In 1978, Zamecnik and Stephenson demonstrated that ASOs can inhibit viral replication *in vitro* ^6, 7^. In 1998, the first ASO drug Fomivirsen received approval for treating cytomegalovirus (CMV) retinitis in immunocompromised patients, demonstrating the feasibility and potential of ASOs as antivirals. Since 2013, eight ASO drugs gained approval to treat a variety of genetic disorders.

Single-stranded RNA targets of ASOs tend to fold by forming intramolecular base pairs, which stack with each other to form double-stranded helices and higher order structures. These structures are often important for biological functions ^8, 9^. These structures often hinder ASO hybridization, as their pairing interactions and 3D geometries are not compatible with each other. We call this the “target RNA folding problem” in ASO designs. Current ASO design strategies either avoid targeting structured regions ^10^ or simply rely on tiling target sequences and screening large numbers of ASOs. The inadequate consideration of structures has resulted in the failure of many ASO candidates and leaves important structured targets as inaccessible. ASOs that have shown therapeutic efficiency, including approved drugs, are often suboptimal in this regard. Their limited target binding affinity and specificity lead to low potency and high toxicity; their high EC_50_ contributes to the challenge of delivering sufficient amounts of drugs to the right tissues and cells.

Coronavirus disease 2019 (COVID-19) is caused by the novel coronavirus severe SARS-CoV-2 ^11, 12^; it has become a serious global pandemic because of its lethality and contagiousness. Variant strains have emerged with enhanced infectivity, even among vaccinated people. Veklury (remdesivir) is the only drug that has received U.S. Food and Drug Administration authorization for treating COVID-19 cases requiring hospitalization ^13^. The world is in dire need for more effective treatments. SARS-CoV-2 is a positive-sense, single-stranded RNA virus with a large genome (close to 30 kb), presenting functionally important sites as drug targets ^14, 15, 16, 17^. The genomic organization of SARS-CoV-2 is highly conserved among viruses in the Nidovirales order, with a large replicase gene occupying the 5′ two thirds of the genome and structural and accessory genes taking up the downstream region ^18^. The replicase gene contains two open reading frames (ORFs), 1a and 1b, which overlap at the frameshifting stimulation element (FSE). FSE directs programmed ribosomal frameshifting (PRF) and is required for ORF1b translation ^19, 20, 21^. As essential steps in their life cycles, these viruses synthesize subgenomic RNAs for expressing downstream genes ^22, 23^; the transcription regulatory sequences (TRS) are central to this process. There have been reports of using ASOs to target PRF and TRS for inhibiting severe acute respiratory syndrome-related coronavirus (SARS-CoV) replication ^16, 24, 25^. Therefore, the well-conserved FSE and TRS have proven to be valid ASO target sites.

In this study, we developed an ASO design method that takes target structures into consideration. Our 3D-ASO design principles may be summarized as compatibility with target structures in 3D space and engineering in tertiary interactions. We identified and validated two design templates and identified potent inhibitors of SARS-CoV-2 by targeting their FSE and TRS regions, respectively.

## RESULTS

### Design templates derived from a pseudoknot structure

We aimed to understand how an ASO and a structured target RNA interact. However, the required structural information is lacking, i.e., few 3D structures of RNA hairpins in complex with ASOs are available. Therefore, we started out by inspecting RNA structures, such as pseudoknots, that resemble the association between ASOs and hairpins. Pseudoknot refers to an RNA hairpin with its loop pairing with an upstream or downstream strand, which may be viewed as equivalent to an ASO (Fig. 1a). We inspected 93 pseudoknot structures (Table S1) and observed extensive tertiary interactions, including major-groove and minor-groove base triples (a third base contacting a base pair from the major- or minor-groove side), non-WC base pairs, and base stacking. Among them, the telomerase RNA pseudoknot structure stands out as having most extensive tertiary interactions (Fig. 1d-f) ^26^.

**Fig. 1.**
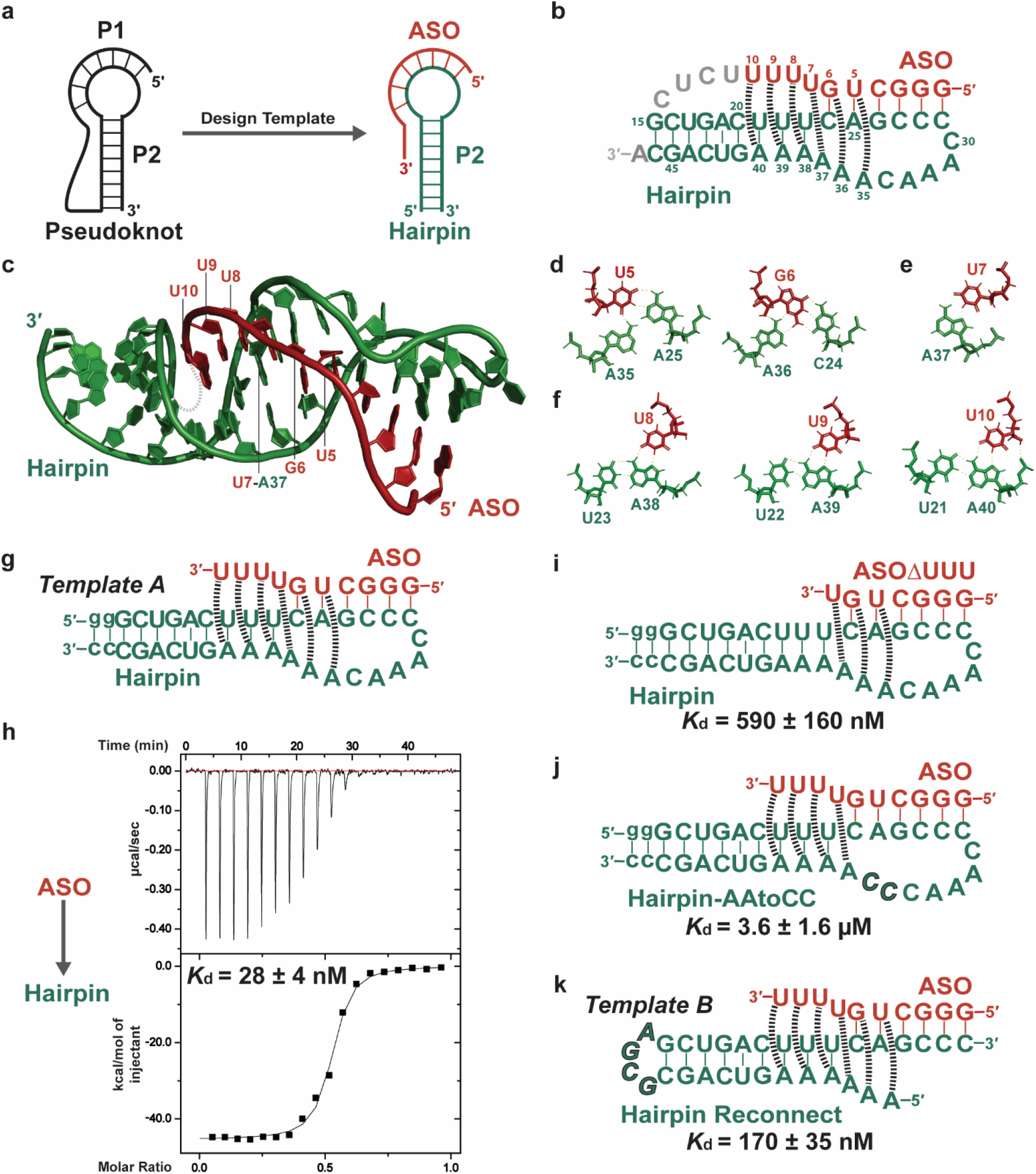
Concept of 3D-ASO design inspired by RNA pseudoknot structures. **a** A pseudoknot may be viewed as an ASO bound to a target hairpin. **b** Secondary structure drawing of the P2b-P3 pseudoknot of human telomerase RNA previously engineered for structural studies ^26^. By omitting the residues shown in silver, we view the structure as a model ASO bound to a Hairpin. Tertiary interactions revealed by the structure are represented by dashed lines and detailed in panels d, e, and f. **c** NMR structures of the pseudoknot (PDB ID: 1YMO) provide a 3D model of ASO-Hairpin complex. The omitted residues are represented by a silver dashed line. **d** Minor-groove base triples formed by U5 and G6 of ASO. **e** A Hoogsteen base pair by U7. **f** Major-groove base triples by U8, U9 and U10. **g** 3D-ASO design template *A*. **h** ITC binding isotherm. Top: raw data. Bottom: the integrated heat values plotted against ASO:Hairpin molar ratio. **i**,**j** Truncation and mutations of the model ASO and Hairpin decrease the affinity, demonstrating the importance of tertiary interactions. *K*_d_ values are shown along with standard errors obtained from fitting. **k** The same ASO may be used to target the basal region of a target hairpin, forming the same set of tertiary interactions. We obtained Hairpin Reconnect by connecting the 5′ and 3′ ends of Hairpin with a GCGA tetraloop and breaking up the Hairpin loop.

We selected several pseudoknot structures, designed ASO-hairpin pairs, and tested their binding with isothermal titration calorimetry (ITC). Indeed, the model ASO-Hairpin pair derived from the telomerase pseudoknot showed high-affinity binding (Fig. 1g,h). Therefore, we used this structure as a template for subsequent biochemical characterization and ASO design work. ASO forms WC pairs with 5′ region of the Hairpin loop using residues G1‒C6 (Fig. 1g). Importantly, according to the telomerase pseudoknot structure, ASO is involved in extensive tertiary interactions. The minor-groove surface of U5 and G6 are contacted by two A residues in the 3′ loop region (A35 and A36 in pseudoknot numbering, Fig. 1d). Three overhanging U residues at the 3′-end of ASO (U8–U10) contacts the base pairs in the Hairpin stem from the major groove side (Fig. 1f). Even though these ASO residues are not involved in WC pairing, they engage in extensive hydrogen bonds and base stacking between consecutive layers of base triples. An important feature of the ASO-Hairpin template is a Hoogsteen pair formed between U7 of the ASO and A37 at the 3′ terminus of the Hairpin loop (Fig. 1e). This Hoogsteen pair is important for proper geometry bridging the minor- and major-groove base triple segments. Overall, these tertiary interactions should contribute to binding affinity and structural specificity.

### Base triple interactions are essential for high-affinity binding

To test if the tertiary interactions observed in the telomerase pseudoknot structure occur in the ASO-Hairpin complex, we disrupted the base triples in ASO-hairpin and measured their affinities using ITC. The *K*_d_ value for ASO/Hairpin is 28 ± 4 nM (Fig. 1h). Truncation of the three U overhanging residues in ASO reduced binding to Hairpin by 20-fold (Fig. 1i). Dramatically, substituting A36 and A37 in the Hairpin loop with two C residues disrupted the ASO binding by 130-fold (Fig. 1j). These results indicate that the major-groove and minor-groove base triples are important in the ASO-Hairpin interaction and therefore suggest that the ASO-Hairpin complex is structurally similar to the telomerase pseudoknot. We call the ASO-Hairpin pair, including its structural and binding features, 3D-ASO design template *A*.

### A 3D-ASO design template that targets hairpin-neighboring regions

ASO binds to the terminal loop of Hairpin (Fig. 1g). Similar sets of tertiary interactions can be engineered into ASOs that hybridize to an RNA strand immediately 3′ to a target hairpin. By simply reconnecting the target RNA strands, we derived Hairpin Reconnect that can similarly bind ASO (Fig. 1k). Indeed, ASO and Hairpin Reconnect do bind but with a 5-fold reduction in affinity, which we attribute to changes in topological constraints. The availability of this derivative design template, which we call template *B*, broadens the applicability of our 3D-ASO design method.

### Base triples and Hoogsteen pairs compatible with the design templates

We surveyed literature and the RNA Base Triple Database ^27, 28, 29^, and identified minor-groove and major-groove base triples structurally compatible with the templates (Fig. 2). These triples have been observed in previously determined RNA structures and therefore are likely to work in 3D-ASO designs. The table of template-compatible base triples helps to determine what resides to use in a 3D-ASO. In template *A* and *B*, the 3′ overhanging residues of ASO form major-groove triples with WC base pairs in the target stem (Fig. 1g,k). For these positions, U works well with A-U and U-A pair, and C is compatible with G-C and C-G pairs (Fig. 2a–d). Additionally, G can form major-groove triples with A-U and G-C pairs, whereas A can form a base triple with C-G (Fig. 2e–g). These purine base triples provide useful alternatives to U and C. Further inward of ASO, U5 and G6 engages form WC pairs with one target residue and interact with another using its sugar surface (Fig. 1d). Any WC base pair can form such a minor-groove triple with A (Fig. 3h–k). It is not clear whether the A can be substituted by another nucleotide. To satisfy the Hoogsteen pair, an ASO can use the WC surface of U to pair with the Hoogsteen surface of an A residue in the target (Fig. 3l). Similarly, G can pair with the Hoogsteen surface of A or G, and protonated C (C+) can pair with G (Fig. 3o).

**Fig. 2.**
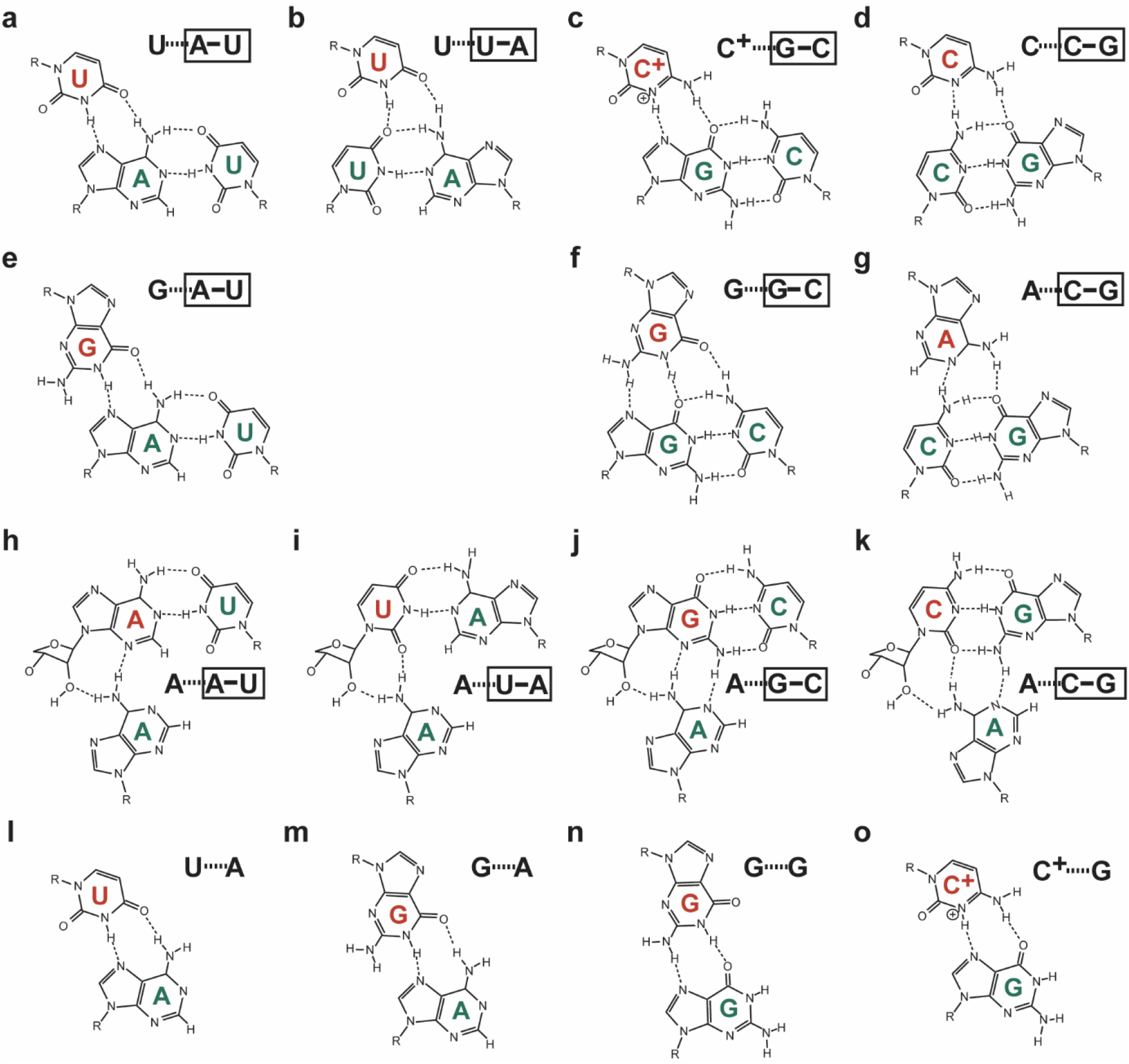
Isosteric tertiary interactions compatible with the design templates. **a**‒**g** Major-groove base triples. **h**–**k** Minor-groove triples. **l**–**o** Hoogsteen pairs that can bridge the segments of major-groove and minor-groove base triples.

**Fig. 3.**
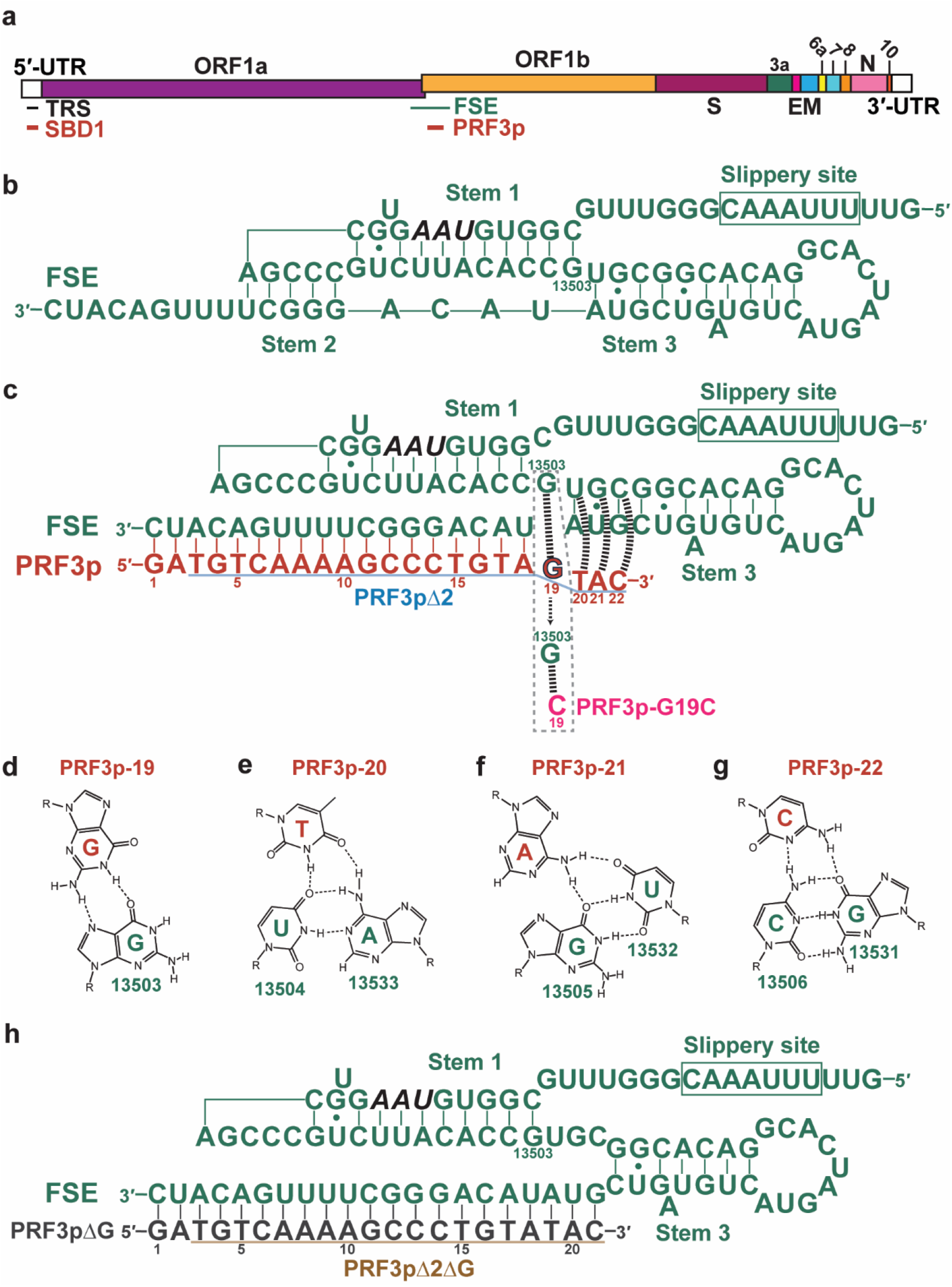
3D-ASO designs targeting the SARS-CoV-2 FSE. **a** Schematic of the genomic RNA of SARS-CoV-2. The TRS and FSE are indicated by horizontal lines. The 3D-ASOs targeting these sites are shown as thick red bars. **b** The pseudoknot structure in FSE causes ribosome stalling at the slippery site (boxed). The UAA stop codon for ORF1a is shown in black italic. **c** PRF3p (an ASO with PMO backbone) is designed to disrupt stem 2 and part of stem 1, allowing the ribosome to proceed to the stop codon and dissociate. Dashed lines represent tertiary interactions. **d** Hoogsteen pair between PRF3p and FSE. **e**–**g** Major-groove base triples between PRF3p and FSE base pairs in Stem 3. **h** Deletion of G19 in PRF3p (PRF3pΔG) results in a 1D-ASO that also disrupts stem 2.

### Design of 3D-ASOs targeting SARS-CoV-2

We designed several ASOs that should be able to forge extensive base-triple and Hoogsteen base-pairing interactions with SARS-CoV-2 FSE and TRS (Fig. 3a). Coronaviruses require FSE to induce programmed –1 ribosomal frameshifting (PRF) and to translate ORF1b, which encodes the RNA-dependent RNA polymerase as the catalytic subunit of the replicase and transcriptase complexes ^20, 30^. FSE contains a pseudoknot structure proceeded by a slippery sequence and an attenuator hairpin (Fig. 3b). During translation, a ribosome is stalled at the slippery site by the pseudoknot and this process gives the ribosome a chance to slip back by one nucleotide before being able to disrupt the pseudoknot and continue translating ^31^. Without frameshifting, the ribosome encounters a stop codon and fall off, producing only protein 1a. With frameshifting, it completes the translation of ORF1b, generating protein 1a-1b.

We designed a 3D-ASO that disrupts the FSE pseudoknot structure with the goal of eliminating the essential PRF for SARS-CoV-2. Cryo-EM and crystal structures of the pseudoknot have been determined ^32, 33, 34^. Despite differing in detail, they agree that the pseudoknot contains stems 1‒3 (Fig. 3b). Our idea is to disrupt stem 2 and a G-C pair in stem 1 so that the ribosome will not stall or will stall at a position downstream the slippery site. The design PRF3p, shown in Fig. 3c, follows template *B*. This 22-nt ASO hybridizes to a region immediately neighboring stem 3 on the 3′ side and engages in a Hoogsteen pair and three major-groove base triples (Fig. 3d–g). We chose the phosphorodiamidate morpholino (PMO) modification because of its exceptional stability, neutral backbone, and safety as demonstrated by approved drugs ^35^.

Previously, Neuman *et al.* designed nine PMO ASOs targeting SARS-CoV RNA and found that the most potent one (named TRS2) inhibits viral replication at 10‒20 μM concentrations by binding to the TRS region ^25^. The authors proposed that the target TRS region folds into a hairpin and designed the 21-nt TRS2 to be complementary to nearly the entire hairpin loop (Fig. 4a). In the two-dimensional (2D) drawing, TRS2 maximizes base pairing with the target without having to disrupt its secondary structure. However, in 3D space the duplex formed by TRS2 and the hairpin loop is relatively rigid, forcing the loop ends to separate by ~56 Å, whereas their immediate neighboring residues in the hairpin stem should be ~11-Å apart (Supplementary Fig. 1). Therefore, either a substantial number of TRS2 bases cannot hybridize to the target loop, or they achieve so with the free-energy expense of having to disrupt the hairpin stem. In either case, TRS2 cannot achieve the intended binding affinity and specificity and any unpaired TRS2 residues only contribute to binding to off-target RNAs.

**Fig. 4.**
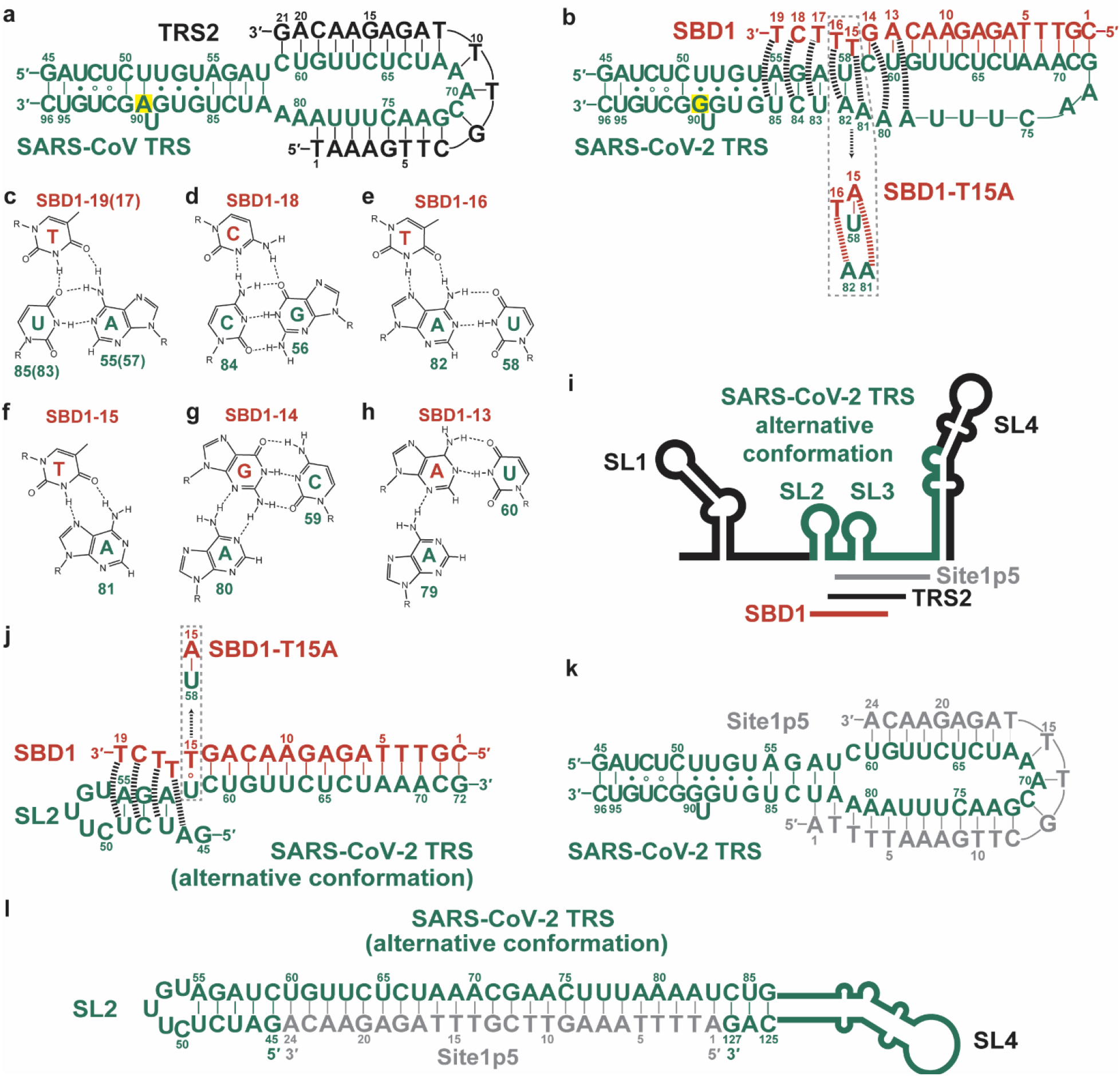
Design of SBD1 targeting the SARS-CoV-2 TRS. **a** A previously reported ASO called TRS2 was proposed to bind to a hairpin loop ^25^. **b** SBD1 with PMO backbone, with a variant T15A shown below. **c**–**h** Designed tertiary interactions between SBD1 and viral RNA. **i**, Recent studies strongly suggest an alternative conformation for the 5′-UTR. The hairpin residues in panel b (colored in green) are involved in three stem-loop structures. **j** SBD1 has to disrupt SL3 but can engage in tertiary interactions with SL2, similar to those shown in panels c, d, and f. **k**,**l** A 2D design (Site1p5) is fully complementary to the target sequence. Panels k and l illustrate how Site1p5 is expected to interact with the TRS region in the single-hairpin and alternative conformations, respectively.

Using template *A*, we designed a 3D-ASO that targets the SARS-CoV-2 TRS, which we call structure-based design 1 (SBD1). The proposed TRS hairpin structure proposed is almost identical between SARS-CoV and SARS-CoV-2, with a single nucleotide substitution (A90G, SARS-CoV-2 numbering) that changes a U-A pair to a U-G wobble pair (Fig. 4a,b). Therefore, the TRS2 ASO should similarly inhibit SARS-CoV and SARS-CoV-2. SBD1 uses 14 nucleotides (shared with TRS2) to form WC base pairs with 5′ region of the loop but leaves nine 3′-loop residues to connect back to the hairpin stem (Fig. 4b). Five residues at the 3′-end of SBD1 (T19, C18, T17, T16, and T15) are designed to form major-groove base triples with the hairpin stem base pairs and a Hoogsteen pair with the 3′ terminal A81 residue of the loop (Fig. 4c–f). The neighboring A80 and A79 residues in the loop can form minor-groove base triples with the SBD1-loop duplex (Fig. 4g,h). SBD1 pairs with three 5′ residues of the core TRS sequence ACGAAC and conformationally constrains the rest. Thus, SBD1 is expected to disrupt the discontinuous transcription.

The design of 3D-ASOs for targeting the TRS region is complicated by alternative secondary structures. In fact, secondary structure prediction and chemical probing analyses strongly suggested that 5′ region of SARS-CoV-2 genomic RNA folds into a structure that contains four stem loop structures (Fig. 4i) ^36, 37, 38, 39, 40, 41, 42^. The sequence proposed to fold into the TRS hairpin occupies SL2, SL3, 5′ region of SL4, and the linkers between them. It is not uncommon for RNAs to fold into alternative structures. We considered both.

SBD1 can forge extensive tertiary interactions with SL2 of the alternative structure (Fig. 4j). SL2 contains a five-base-pair stem and a pentaloop. To bind SBD1 via template *A*, two base pairs at the base of SL2, G45-C59 and A46-U58, are replaced by interactions with SBD1, including a Hoogsteen pair between A46 and SBD1-T16, a non-canonical pair between U58 and SBD1-T15, and WC pair between C59 and SBD1-G14. Importantly, three residues at the 3′-end of SBD1 (T19, C18, and T17) can form major-groove base triples with the SL2 stem and recognize the structure, resembling the expected interactions with the TRS hairpin (Fig. 4b,j). No strong minor-groove interactions are expected. SL3 contains only four base pairs with a heptaloop and is not expected to be very stable. SBD1 is designed to disrupt SL3 via WC base pairing. Remarkably, despite having to disrupt part of SL2 and engaging in fewer tertiary interactions with the alternative structure, SBD1 is expected to bind both potential structures.

For comparison, we designed Site1p5 that forms WC pairs throughout the 24-nt region between SL2 and SL4, occupying the TRS core and surrounding regions (Fig. 4k,l). Like SBD1, Site1p5 disrupts SL3. The duplex formed by Site1p5 and the target strand should stack with the SL2 and SL4 stems, but otherwise no tertiary interactions are expected. Thus, Site1p5 is a 2D design.

### Identification of two lead ASOs that inhibit SARS-CoV-2

We generated synthetic ASOs with PMO backbone and tested their abilities to inhibit SARS-CoV-2 replication in cultured HEK293 cells expressing the human angiotensin-converting enzyme 2 (hACE2) receptor ^43^. We first transfected HEK293-hACE2 cells with ASOs at 10 and 20 μM concentrations. Viral inoculum was added the following day. At 24 h post infection (hpi), viral loads were examined using immunocytochemistry (ICC) to detect the Spike protein and qRT-PCR to measure viral RNAs. Consistent with the previous report, TRS2 showed limited inhibition of the virus compared to the mock transfection (without ASO) (Fig. 5a) ^25^. Our 2D design, Site1p5, showed little inhibition of the virus. In contrast, both 3D-ASOs, SBD1 and PRF3p, showed strong inhibition, with barely any viral signals detected in the latter. These visual results were corroborated by quantifying the number of spike-positive infected cells (Fig. 5b).

**Fig. 5.**
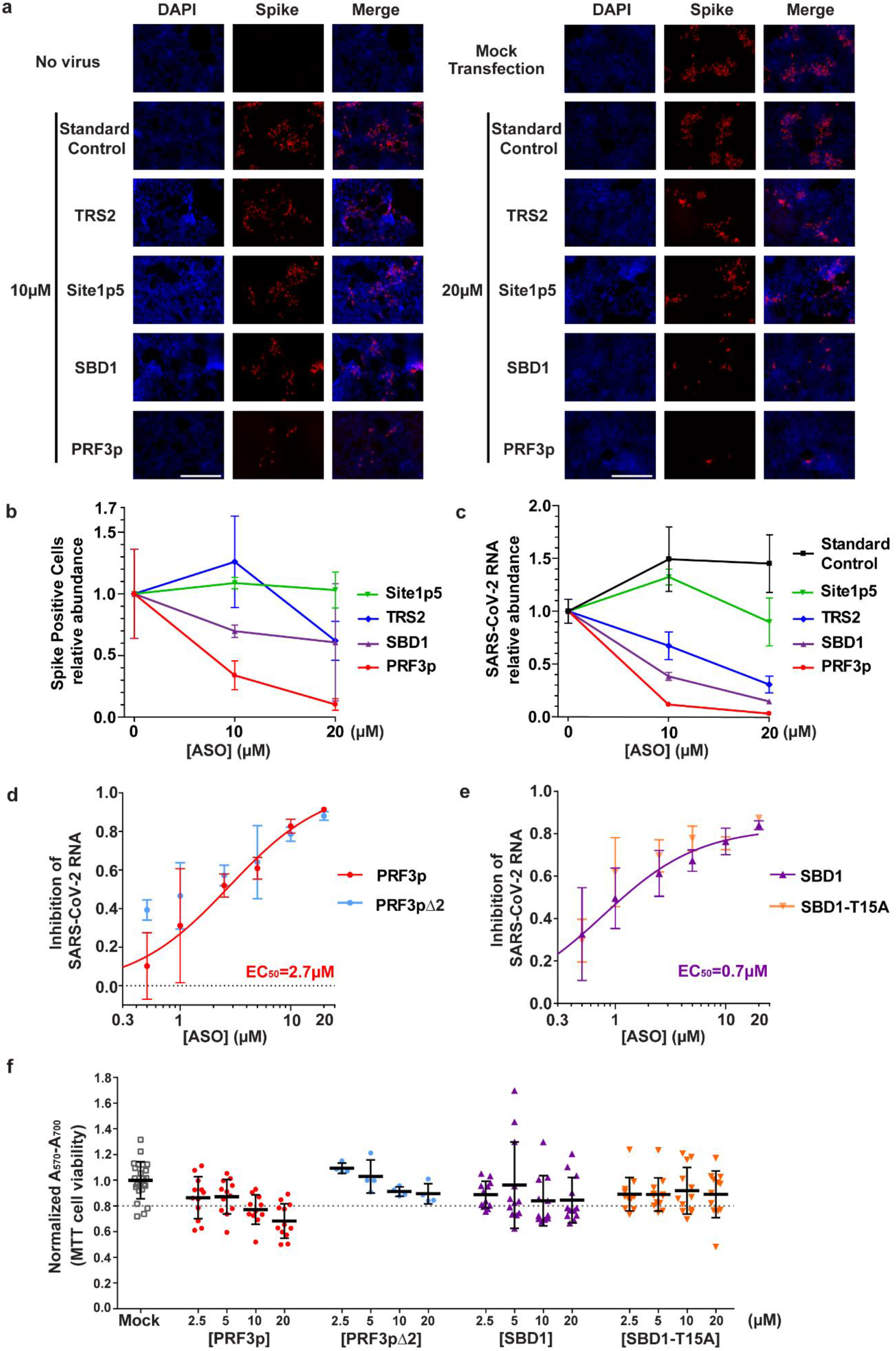
Identification of lead 3D-ASO inhibitors. **a**–**e** HEK293-hACE2 cells was infected by the SARS-CoV-2 virus. ICC and total RNA extraction were performed at 24 hpi. **a** ICC images of cells transfected with ASOs at 10 and 20 μM prior to infection, stained for the spike antigen. The scale bars represent 200 μm. **b** Quantification of spike-positive cells. The data shown are average ± SD. **c** qRT-PCR quantification of viral RNA (a N-coding sequence). The data were normalized to host *GAPDH* mRNA. Shown are average ± SD from three (panel c) or four (panels d, e, f) independent repeats. **d**,**e** Dose-dependent inhibition by PRF3p and SBD1, respectively, along with their variants. **f** Scatter plots of MTT cell viability assays. Horizontal bars indicate means and error bars represent SD.

The qRT-PCR data confirmed the SARS-CoV-2 inhibitory activity of our 3D-ASOs. At 10 and 20 μM, PRF3p strongly inhibited SARS-CoV-2 replication, reducing 86% (*P* = 0.0001) and 96% (*P* = 0.00002) of viral RNA (normalized by a host RNA) relatively to that in the mock transfection (Fig. 5c). SBD1 reduced the viral RNA by 56% (10 μM, *P* = 0.0005) and 83% (20 μM, *P* = 0.00007). These results are highly significant as indicated by the *P* values. The data confirmed the mild inhibitory activity of TRS2 (23% at 10 μM, *P* = 0.009; 64% at 20 μM, *P* = 0.001) and the lack of activity for Site1p5. Overall, the ICC and qRT-PCR results demonstrate that our structure-based designs are superior at inhibiting viral replication relative to the conventional designs.

We next examined dosage dependency of inhibition by the 3D-ASOs. SARS-CoV-2 viral inhibition assays were performed with 3D-ASO concentrations from 0.5 to 20 μM. The abundance of viral RNAs was measured using qRT-PCR. The data for PRF3p and SBD1 were fit to a dose-dependent curve, resulting in EC_50_ values of 2.7 and 0.7 μM, respectively (Fig. 5d,e). The fitting of the PRF3p data expanded the 0–1 inhibition range, whereas that of SBD1 only plateaued to ~86%. In addition, SBD1 appeared to inhibit stronger at concentrations at or below 1 μM.

### The antiviral activity of 3D-ASOs is not due to cytotoxicity

To evaluate their potential cytotoxicity, we transfected HEK293 cells with the ASOs and measured their metabolic activity as a proxy for cell viability using MTT assays. SBD1 is not toxic to cells at all concentrations tested, with normalized viability ranging from 85% to 96%. With 2.5 and 5 μM of PRF3p, cell viability was maintained at around 87%, also not toxic (Fig. 5f). At 10 and 20 μM, PRF3p reduced cell viability to 77% ± 11% and 68% ± 13%, respectively, suggesting that at these PRF3p concentrations the viral inhibition may be compounded by cytotoxicity.

To identify the origin of the PRF3p cytotoxicity at high doses, we searched the human genome and found that 15 nucleotides at its 5′ end are complementary to the mRNA encoding glucose-6-phosphate isomerase (GPI) as the only significant hit. GPI is essential for both catabolic glycolysis and anabolic gluconeogenesis. We truncated two nucleotides from the 5′ end (PRF3pΔ2), transfected it to HEK293 cells, and performed MTT assays. The viability is around 90% at both 10 and 20 μM, not significantly different from that of the mock transfection. Therefore, by deleting two nucleotides from 5′ end, we eliminated the slight cytotoxicity of PRF3p. We tested PRF3pΔ2 using cellular SARS-CoV-2 viral inhibition assay and found its antiviral activity is similar to that of PRF3p (Fig. 5d). Therefore, we conclude that the antiviral activities of 3D-ASOs are likely due to direct interaction with viral RNAs.

### The Hoogsteen-pairing residue of PRF3p is important for antiviral activities

The Hoogsteen pair bridging the major- and minor-groove base-triple segments is an essential component of our 3D-ASO design templates. G19 of PRF3p uses its WC surface to hydrogen-bond with the Hoogsteen surface of G13503 at the 3′ end of stem 1 (Fig. 3b,c,d). Deletion of G19 (ΔG) is expected to disrupt major-groove triples involving residues T20, A21, and C22. Instead, ΔG results in an ASO that is perfectly complementary to the target sequence and is also expected to disrupt stem 2 of the FSE pseudoknot (Fig. 3h). In this sense, the ΔG variants represent the closest matching 1D designs of our 3D-ASOs.

We compared PRF3p and PRF3pΔ2 with their respective ΔG variants using viral inhibition assays. The PRF3p and PRF3pΔG data largely overlap (Fig. 6a). The only statistically significant differences occurred at 10 and 20 μM, where PRF3p inhibits at greater extents (83% and 91%) than PRF3pΔG (86% and 73%), with *P* values of 0.02 and 0.04 respectively. The PRF3pΔ2 and PRF3pΔ2ΔG data also largely overlap, but with statistically significant differences occurring at 0.5 and 2.5 μM (Fig. 6b). At these concentrations, PRF3pΔ2 inhibits viral replication to greater extents (39% and 57%) than PRF3pΔ2ΔG (15% and 49%), with P values of 0.004 and 0.02 respectively. These data suggest that our PRF inhibitor 3D-ASOs are stronger than their closest 1D counterparts.

**Fig. 6.**
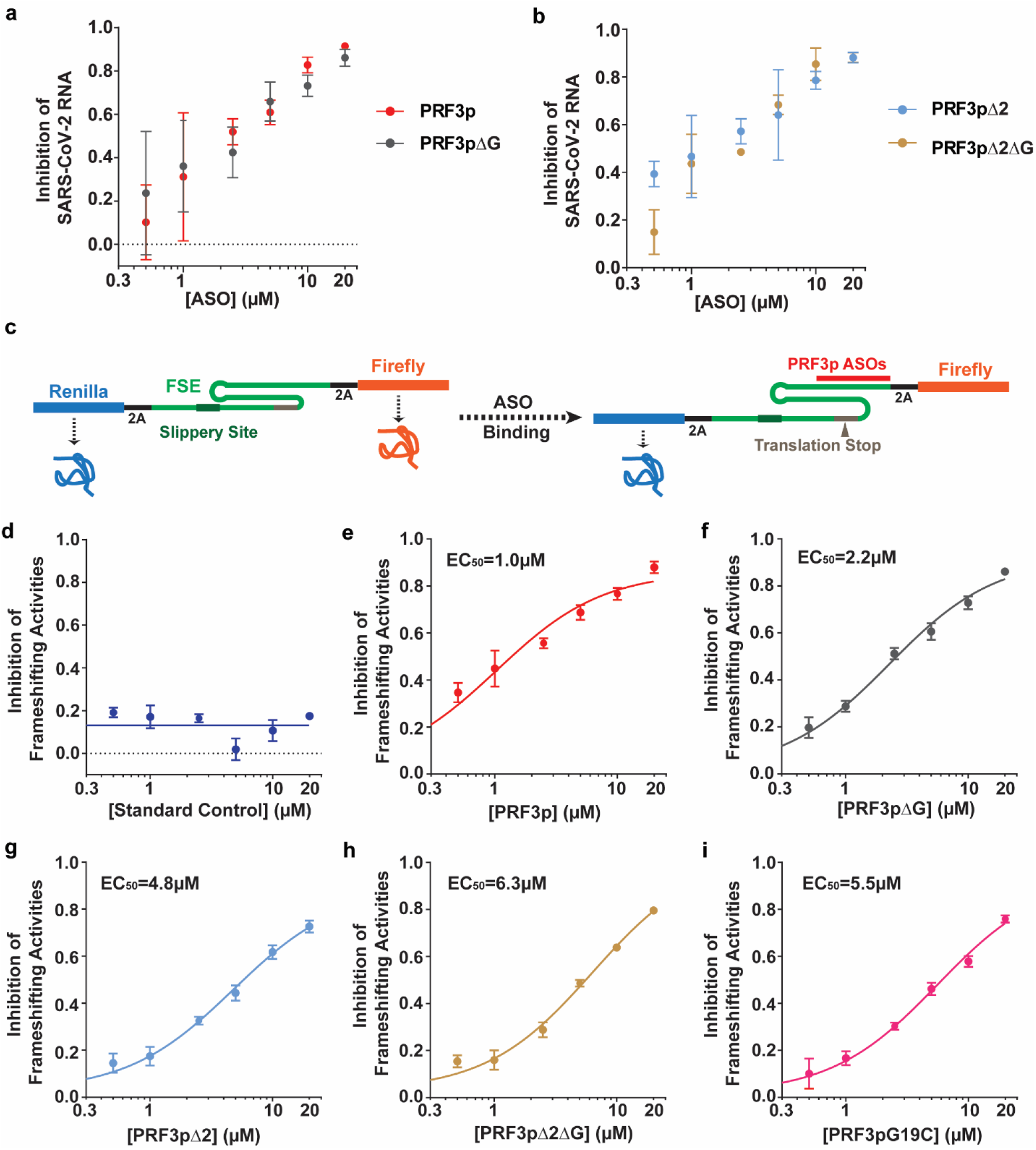
Hoogsteen-pairing residue of PRF3p is important for activity. **a**,**b** Cellular SARS-CoV-2 viral inhibition assays comparing PRF3p and PRF3pΔG (a), and PRF3pΔ2 and PRF3pΔ2ΔG (b). **c**–**i** Test PRF3p and its variants using the dual luciferase PRF reporter assay and HEK293 cells. **c** Illustration of the CoV2-FSE reporter construct. PRF3p is designed to bind and disrupt the pseudoknot structure of FSE, interrupt recoding at the slippery site, and cause translation to stop. 2A, also called StopGo, sequences reprogram translation to produce isolated Firefly and Renilla luciferases rather than fusion proteins. **d**–**i** PRF reporter activities of indicated ASOs. The data plotted are average ± SD from 4 repeats.

To more accurately compare the PRF3p variants, we constructed a dual luciferase-based reporter for measuring their PRF inhibitory activities. In this reporter, the SARS-CoV-2 FSE was inserted between the coding sequences for Renilla and Firefly luciferases, which are equivalent to the 1a and 1b viral proteins (Fig. 6c). To prevent protein fusion from interfering with luciferase activities and stabilities, two StopGo sequences were inserted to flank the FSE ^44^. Whereas the Standard Control ASO did not alter the frameshifting activity, all PRF3p variants induced dosage-dependent inhibition (Fig. 6d–h), clearly demonstrating that the latter inhibit SARS-CoV-2 replication by inhibiting the PRF activities of the FSE. Curve fitting yielded EC_50_ values of 1.0 μM (95% CI: 0.7–1.5 μM) for PRF3p and 2.3 μM (95% CI: 1.8––2.8 μM) for PRF3pΔG (Fig. 6e,f). The increase in EC_50_ caused by ΔG is statistically significant because their 95% CI ranges do not overlap. The EC_50_ for PRF3pΔ2 (4.8 μM, 95% CI: 3.7–6.4 μM) is also lower than its 1D counterpart 3pΔ2ΔG (6.4 μM, 95% CI: 4.9–8.3 μM) (Fig. 6g,h). A protonated C should also be able to mediate the Hoogsteen pair instead of G19 in PRF3p (Fig. 2o). However, the protonation requirement makes the interaction relatively weak at physiological pH. We compared PRF3p and its G19C mutant using the PRF reporter assay. The EC_50_ of G19C is 5.5 μM, 5.5 folds that of PRF3p (Fig. 6i). These results highlight the importance of the Hoogsteen pair in 3D-ASO design.

### A variant of SBD1 inhibits SARS-CoV-2 slightly better

We designed a variant of SBD1 (T15A) that optimizes the interactions with the alternative TRS conformation (Fig. 4j). The T15A mutation changes a T-U58 non-canonical pair to an A-U58 pair. SBD1-T15A can engage in alternative interactions with the TRS region in the single-hairpin conformation. A15 of SBD1-T15A cannot form a Hoogsteen pair with target A81. Instead, A15 can form a WC pair with U58, disrupting its pairing with A82 (Fig. 4b). This switching of pairing partner of U58 converts the major-groove base triple between T16 and U58-A82 to a Hoogsteen pair between T16 and A82. We expect A81 to form a minor-groove base triple with the A15-U58 pair, compensating the loss of the major-groove triple. In cellular SARS-CoV-2 inhibition assays, SBD1-T15A inhibits slightly better than SBD1 at most concentrations (Fig. 5e). The increases are statistically significant, with *P* value of 0.03 at both 5 and 20 μM. Like SBD1, T15A is not toxic to cells (Fig. 5f).

## DISCUSSION

Our study demonstrates that 3D-ASOs are better than conventional designs. SBD1 inhibits SARS-CoV-2 much better than TRS2 and Site1p5, with which it shares a stretch of 13-nucleotide sequence. Yet the 19-nt SBD1 is shorter than the 21-nt TRS2 and 24-nt Site1p5. The PRF3p and PRF3pΔ2 also inhibit SARS-CoV-2 and PRF better than their closest matching 1D ASOs (the corresponding ΔG variants). The improvements are not as substantial as what SBD1 achieved over TRS2 and Site1p5. This may be explained by the fewer tertiary interactions engineered and by the fact that the target RNA adopts different conformations when bound to the PRF3p 3D and 1D ASOs (Fig. 3c,h).

3D-ASO design requires understanding of target RNA structures. This understanding can be obtained via secondary structure prediction, structural probing, and 3D structure determination. RNA secondary structure prediction parameters and algorithms have been developed for decades ^45^. Strong RNA secondary structures can often be predicted with high confidence. Such prediction can be performed on any target RNA sequences. Additionally, structural probing can be performed both *in vitro* and in cells, and high-throughput methods are available ^46, 47^. The results can be used as experimental constraints to improve the accuracy of secondary structure predictions. 3D structure determination of RNAs has been challenging but new methods are being developed ^48, 49, 50^. We should also keep in mind that RNA structures can be dynamic and conformational changes are often required for their biological functions.

When performing 3D-ASO design, it is important to distinguish the ASO-free and bound states of a target RNA. The design templates presented here represent the bound state, whereas the RNA structure prediction and characterization often concern the ASO-free state. In some cases, such as SBD1 binding to the TRS hairpin (Fig. 4b), we assume the target RNA structure is largely unchanged to maximally facilitate ASO binding. Even though recent reports strongly suggest an alternative conformation for the ASO-free TRS region (Fig. 4i), the single-hairpin conformation can still be thermodynamically stable when bound to SBD1 (Fig. 4b). In fact, SBD1 can bind to both conformations of TRS via template *A* and *B* (Fig. 4b,j), respectively.

To fully unleash the power of 3D-ASO design, chemical modifications need further exploration to determine their compatibility with the design templates. Backbone modifications are required for increasing ASO *in vivo* stability, enhancing target affinity, and reducing toxicity. This study shows that PMO is compatible with the 3D-ASO templates but how the PMO backbone affects the tertiary interactions remains to be determined. Phosphorothioate does not dramatically change nucleic acid geometry and only introduces subtle changes in backbone geometry; therefore, we expect it to be well-tolerated at all 3D-ASO positions. Compared with native RNAs, ribose modifications such as 2′-O-methoxyethyl (2′-MOE), 2′-O-methyl (2′-OMe), and locked nucleic acid (LNA) ^51^ make the 2′-functional group bulkier. They should be tolerated at most positions in the 3D-ASO template structure, in which 2′-OH are open to the solvent. At positions 4–7, the riboses are in relatively crowded surroundings (Fig. 1c) and therefore 2′-fluoro may be used instead. Like PMO, peptide nucleic acid (PNA) ^52^ ASOs are very resistant to both nuclease and protease degradation and has a neutral backbone. PNA and complementary RNAs hybridize to adopt an A-form helical conformation, with the PNA chain being more flexible ^53, 54^. Therefore, PNAs should fit into our design templates. Further, base modifications can be used to enhance the tertiary interactions. The knowledge from exploring these modifications to be readily generalizable to guide 3D-ASO development.

Our structure-based design strategy has the potential to substantially improve the therapeutic efficacies of ASOs. Although current FDA-approved ASOs are 18–30 nucleotides long, our study suggests that shorter ASOs may be able to achieve sufficient affinity and specificity by engaging in tertiary interactions with targets. A decrease in ASO size would allow more efficient delivery into cells, more favorable bio-distribution, and lower manufacturing costs.

There have been efforts to develop ASO and other synthetic oligonucleotide drugs targeting viruses that cause respiratory diseases ^55^. Viruses that have been targeted previously include SARS-CoV ^16, 24, 25^, influenza A ^56^, respiratory syncytial virus (RSV) ^57^, adenovirus, and most recently SARS-CoV-2 ^41, 58^. The ASOs reported in the latter studies inhibit SARS-CoV-2 replication in cultured cells at sub-micromolar concentrations. Because they use modifications (2′-MOE and LNA, respectively) different from PMO used here, their EC_50_ values are not directly comparable. These ASOs hybridize to viral RNA sites that do not overlap with the 3D-ASOs reported here, including the 5′- and 3′-UTRs and coding sequences, and thus are great candidates to apply 3D design for further improvements. A small interfering RNA (siRNA) RSV01 was developed by Alsylam for treating RSV infections in lung transplant patients ^57^. Encouragingly, their phase IIb trial showed significant efficacy in preventing RSV infections in 3–6 months. In addition, the siRNA is delivered via nasal sprays, making it very easy to administer. As siRNAs are also short synthetic oligonucleotides, our 3D-ASO drug candidates may be administered either as a nasal spray or intravenously. The convenience of the nasal spray formulation is particularly attractive as it will make the drug more accessible to underserved/disadvantaged populations and allow administration outside of the controlled healthcare settings required for IV delivery.

## METHODS

### Synthetic oligos

The following RNA oligos were synthesized by Sigma:

ASO: 5′-GGGCUGUUUU-3′
ASOΔUUU: 5′-GGGCUGU-3′
Hairpin Reconnect: 5′-AAAAAAGUCAGCGCGAGCUGACUUUCAGCCC-3′

The following DNA oligos were synthesized by Integrated DNA Technologies:

Hairpin transcription template strand 5′-GGGCTGACTTTTTTGTTTGGGGCTGAAAGTCAGCCC<u>TATAGTGAGTCGTATTA</u>-3′
Hairpin AAtoCC transcription template strand 5′-GGGCTGACTTTTGGGTTTGGGGCTGAAAGTCAGCCC<u>TATAGTGAGTCGTATTA</u>-3′
T7 promoter sense strand 5′-<u>TAATACGACTCACTATA</u>GG-3′ (The T7 promoter sequence is underlined.)
2019-nCoV_N1-F 5′-GACCCCAAAATCAGCGAAAT-3′
2019-nCoV_N1-R 5′-TCTGGTTACTGCCAGTTGAATCTG-3′
Human GAPDH qPCR forward 5′-CCACCTTTGACGCTGGG-3′
Human GAPDH qPCR reverse 5′-CATACCAGGAAATGAGCTTGACA-3′
SARS-CoV-2-FSE-forward 5′-CCC<u>AAGCTT</u>ACCCATGCTTCAGTCAGCTG-3′ (The HindIII site underlined)
SARS-CoV-2-FSE-reverse 5′-GGA<u>AGATCT</u>GTCAAAAGCCCTGTATACGAC-3′ (The BglII site underlined)
PMO oligos (Table 1) were synthesized by Gene Tools.

**Table 1.**
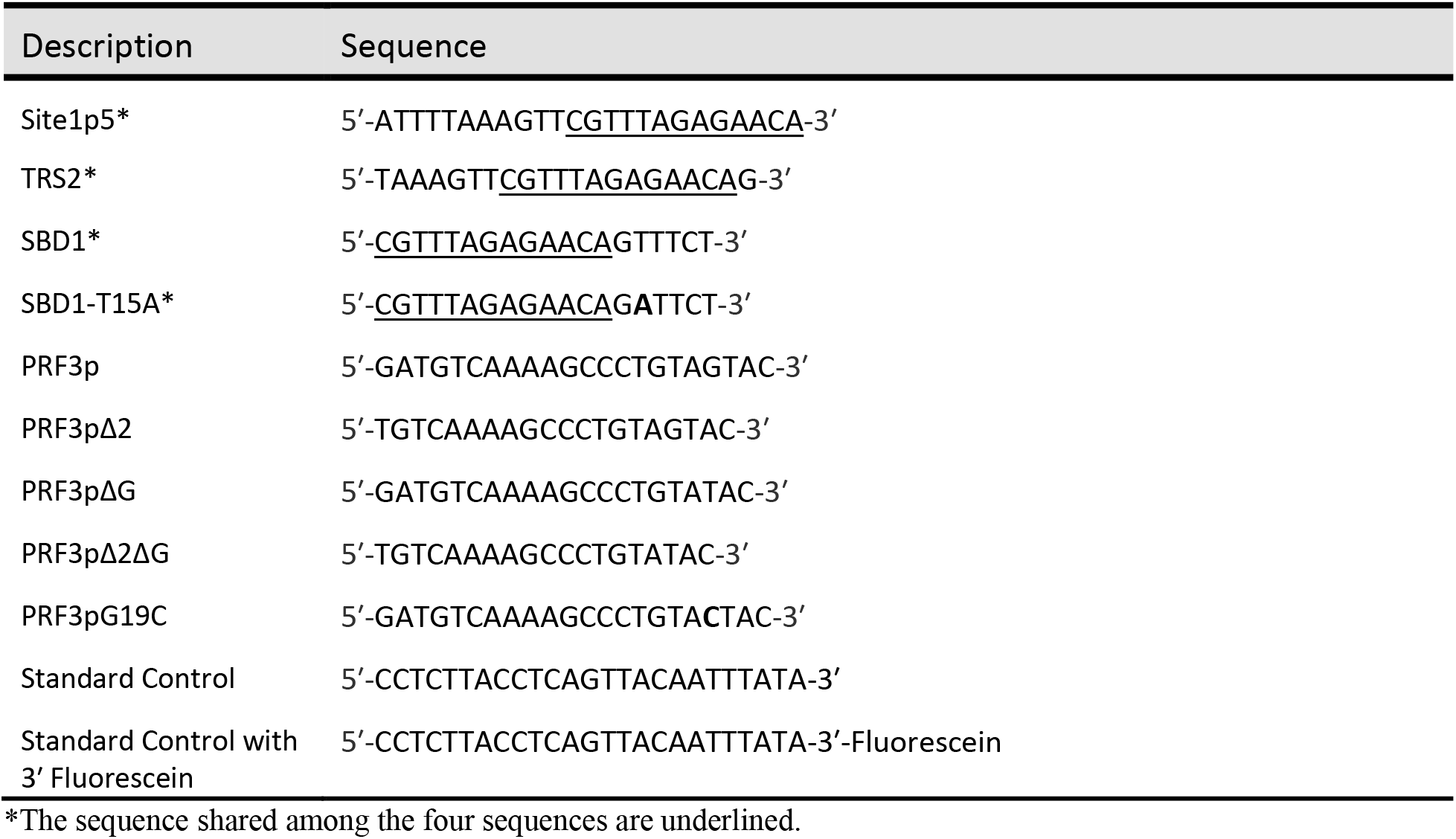
PMO ASO sequences used.

### Transcription and purification of RNAs

The Hairpin and its AAtoCC mutant RNAs were transcribed *in vitro* using T7 RNA polymerase and synthetic DNA templates as described ^50^. Briefly, the template strands were mixed with the T7 promoter sense strand equimolarly to generate the transcription templates. The transcription reactions were set up at a 5-mL volume, ethanol-precipitated and purified using denaturing tube gels. In these gels, RNA bands were separated via electrophoresis on a 491 Prep Cell (BIO-RAD) and were collected with an AKTA Prime chromatography system (Cytiva). The RNAs were concentrated and buffer-exchanged into 10 mM NaHEPES pH 7.0 using Amicon^®^ Ultra-15 centrifugal filter units with 3-kDa molecular weight cutoff (Millipore).

### Construction of the CoV2-FSE119 reporter plasmid

A 119-bp SARS-CoV-2 FSE sequence was synthesized as a gBlock by Integrated DNA Technology and was amplified using primers SARS-CoV-2-FSE-forward and SARS-CoV-2-FSE-reverse. The PCR product was digested by HindIII and BlgII and ligated into pSGDLuc 3.0 44.

### Isothermal titration calorimetry

ITC was performed using a Microcal iTC-200 isothermal titration calorimeter (GE Healthcare). The binding buffer contained 2 mM NaHEPES pH 7.0 and 100 mM KCl. In all experiments, 100 μM ASO was titrated to 200 μL of 10 μM target hairpin. The data were analyzed using Origin 7.0 (Microcal).

### Cellular SARS-CoV-2 viral inhibition assays

All experiments involving live viruses were conducted in a UCLA BSL3 high-containment facility. We has previously established cellular models for SARS-CoV-2 infection ^43^. SARS-CoV-2 (isolate USA-WA1/2020) was obtained from BEI Resources of NIAID. SARS-CoV-2 was passaged once in Vero E6 cells and viral stocks were aliquoted and stored at −80°C. HEK293-hACE2 cells were cultured in 96-well plate at 37°C in 5% CO2. Each well was seeded with ~5,000 cells in 100 μL media containing DMEM (Thermo Fisher) with 10% FBS and 0.01% puromycin. Cells were transfected by incubation in fresh media with ASOs at specified concentrations and mixed well with the Endo-Porter PEG transfection reagent (Gene Tools, 0.6 μL/100 μL media). The following day, the transfected cells was inoculated with the virus at a multiplicity of infection of 0.1. At 24 hpi, cells were either fixed to inactivate the virus and viral loads were examined using ICC and qRT-PCR.

### Immunocytochemistry

Cells were fixed with methanol to inactivate the viruses and were incubated in −20°C freezer until washed with 1x phosphate-buffered saline (PBS) for 30‒60 min at room temperature (RT). Cells were washed for a total of 3 times with 1x PBS and permeabilized using blocking buffer (2% BSA, 0.3% Triton X-100, 5% Donkey Serum, 5% Goat Serum in 1x PBS) for 1 h at RT. For immunostaining, cells were incubated overnight with the primary antibody (Monoclonal Anti-SARS-CoV S Protein (similar to 240C), NR-616 from the BEI Resources, NIAID, NIH) at 4°C. The cells were then washed three times with 1x PBS and incubated for 1 h at RT with respective secondary antibody. DAPI (4′,6-diamidino-2-phenylindole, dihydrochloride) (Thermo Fisher) at a dilution of 1:5,000 in 1x PBS was used to stain cell nuclei. The images shown in Figs. 5a and 5b were acquired at 20× magnification using an Axio Observer D1 inverted fluorescent microscope (ZEISS).

### Genome quantification by qRT-PCR

Total RNAs were extracted using either RNeasy Mini or RNeasy 96 kit (QIAGEN). Their cDNAs were generated using the High Capacity cDNA Reverse Transcription Kit (Applied Biosystems). Quantitative PCR reactions were performed to amplify the N gene of SARS-CoV-2 and the host GAPDH gene using primers listed above, the Platinum SYBR Green qPCR SuperMix-UDG w/ROX kit, and a QuantStudio 3 real-time PCR system (Thermo Fisher). The data were analyzed using the Thermo Fisher Connect Platform. The dose-dependent data in Fig. 5c–e were fit to the equation 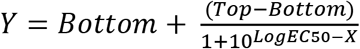 using PRISM (version 7, GraphPad).

### MTT cell viability assays

HEK293 cells were seeded at 5,000 cells per in 96-well plates and transfected as described above. Forty-eight hours posttransfection, the cells were stained and solubilized using the CellTiter 96 Non-Radioactive Cell Proliferation Assay Kit (Promega). The plates were shaken to obtain uniformly colored solutions. The absorbances at 570 nm were recorded at SpectraMax M5 microplate reader (Molecular Devices). The reference wavelength was set to 700 nm to reduce background.

### Programmed ribosomal frameshifting reporter assays

HEK293 cells were seeded at 5,000 cells per well in 96-well plates. When the cells reached about 25% confluency, we transfected them with ASOs using Endo-Porter. Sixteen hours after ASO transfection, when cells usually reached confluency of around 70%, the CoV2-FSE119 plasmid was transfected using lipofectamine 3000 (Thermo Fisher Scientific) at 100 ng per well. Forty-eight hours after plasmid transfection, cells were lysed, and the luciferase activities were measured using the Dual-Luciferase^®^ Reporter Assay System (Promega). Signals were recorded using SpectraMax M5 microplate reader (Molecular Devices) with a 2-sec premeasurement delay followed by a 10-sec measurement. Inhibition of frameshifting activities is calculated as 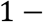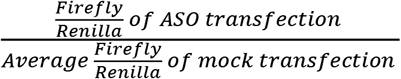. Mean and SD were calculated based on 4 independent experiments. The dose-dependent data in Fig. 6e–i were fit to the equation 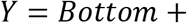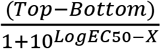 using PRISM.

### Statistical analyses

The Student t-tests, two-tailed and two-sample equal variance, were performed using Excel (Microsoft).

## Data availability

All data are available in the main text or the supplementary information.

## Acknowledgments

We thank Arjit Jeyachandran and Martin Phillips for technical support, Craig L. Zirbel and Neocles Basil Leontis for using the RNA Base Triple Database, Doug Black and Andrey Damianov for advice on computational search for potential off targets, and Gary Loughran and John F. Atkins for advice on the dual luciferase reporter. The following reagent was obtained through BEI Resources, NIAID, NIH: Monoclonal Anti-SARS-CoV S Protein (Similar to 240C), NR-616.

## Author contributions

F.G. conceived the project. Y.L. inspected the structures and performed the biochemical analyses, the PRF reporter assays, and the MTT and qRT-PCR experiments. V.A. and G.G. developed the cellular SARS-CoV-2 viral inhibition assays and did the infection experiments. F.G. and Y.L. wrote the paper with input from other authors.

