## Supplementary information for "Structure-based design of antisense oligonucleotides that inhibit SARS-CoV-2 replication"

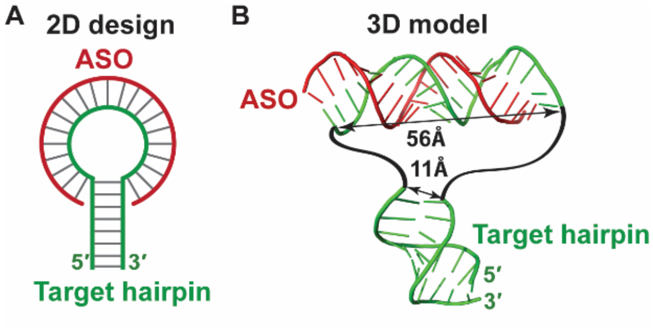


**Supplementary Figure 1. An example to explain why it is important to perform 3D-ASO design. a** A 2D design in which a 21-nt ASO is fully complementary to the entire hairpin loop of a target RNA. **b** 3D model indicates that the ASO-loop duplex and the hairpin stem are incompatible because immediately neighboring residues are far apart. The impossible linkages are indicated by the curved black lines.

**Supplementary Table 1. PDB codes of pseudoknot structures inspected in this study.**

| | | | | | | | | | |
|---|---|---|---|---|---|---|---|---|---|
| 2LC8, | 1F27, | 1L2X, | 1L3D, | 2G1W, | 3T4B, | 5KMZ, | 1E95, | 1KPY, | 1KPZ |
| 1RNK, | 1YG4, | 1YMO, | 2A43, | 2AP5, | 2K95, | 2RP1, | 2RP0, | 2TPK, | 1A60 |
| 1HVU, | 1KAJ, | 1KPD, | 1YG3, | 2AP0, | 2K96, | 2M8K, | 437D, | 4ATO, | 3SUH |
| 3SUX, | 3SUY, | 5K7C, | 5K7D, | 5K7E, | 3FU2, | 3K1V, | 5TPY, | 3VRS, | 4EN5 |
| 4ENA, | 4ENB, | 4ENC, | 2H0S, | 2H0W, | 2H0X, | 2H0Z, | 2N8V, | 4OJI, | 4RGF |
| 2GCS, | 2GCV, | 2HO6, | 2HO7, | 2L1V, | 3MJ3, | 3MJB, | 4JF2, | 2MIY, | 3GX2 |
| 4OQU, | 4P5J, | 4RGE, | 5D5L, | 6FZ0, | 3GX3, | 3GX5, | 3GX6, | 3GX7, | 5KH8 |
| 5NWQ, | 5NY8, | 5NZ3, | 5NZ6, | 5NZD, | 5O62, | 5O69, | 2QWY, | 3Q50, | 3Q51 |
| 4RMO, | 4RZD, | 4LVV, | 4LVW, | 4LVX, | 4LVY, | 4LVZ, | 4LW0, | 2VAZ, | 4PQV |
| 1SLS, | 2IL9, | 5KQE | | | | | | | |
